# *Neisseria gonorrhoeae* co-opts C4b-binding protein to enhance complement-independent survival from neutrophils

**DOI:** 10.1101/2022.12.08.519701

**Authors:** Lacie M. Werner, Allison Alcott, Frida Mohlin, Jocelyn C. Ray, Meagan Belcher Dufrisne, Asya Smirnov, Linda Columbus, Anna M. Blom, Alison K. Criss

## Abstract

*Neisseria gonorrhoeae* (Gc) is a human-specific pathogen that causes the sexually transmitted infection gonorrhea. Gc survives in neutrophil-rich gonorrheal secretions, and recovered bacteria predominantly express phase-variable, surface-expressed opacity-associated (Opa) proteins (Opa+). However, expression of Opa proteins like OpaD decreases Gc survival when exposed to human neutrophils *ex vivo*. Here, we made the unexpected observation that incubation with normal human serum, which is found in inflamed mucosal secretions, enhances survival of Opa+ Gc from primary human neutrophils. We directly linked this phenomenon to a novel complement-independent function for C4b-binding protein (C4BP). When bound to the bacteria, C4BP was necessary and sufficient to suppress Gc-induced neutrophil reactive oxygen species production and prevent neutrophil phagocytosis of Opa+ Gc. This research identifies for the first time a complement-independent role for C4BP in enhancing the survival of a pathogenic bacterium from phagocytes, thereby revealing how Gc exploits inflammatory conditions to persist at human mucosal surfaces.

**Author Summary:** Gonorrhea is considered an urgent threat to public health with an estimated 98 million cases occurring annually worldwide, growing antimicrobial resistance, and the absence of a gonococcal vaccine. Currently, we do not understand how *N. gonorrhoeae* expressing opacity (Opa) proteins survive neutrophil defenses and are recovered viable from infected patients. Here, we investigated how soluble elements of gonorrhea infection, present in human serum, contribute to *N. gonorrhoeae* survival from neutrophils. We found that the serum component C4b-binding protein (C4BP) protects *N. gonorrhoeae* from neutrophil killing and suppresses neutrophil activation. C4BP limited neutrophil phagocytosis of *N. gonorrhoeae* that expressed Opa proteins that bound to neutrophil receptors of the CEACAM family. This work provides novel insight into the interplay between the noncellular and cellular aspects of the innate immune response to *N. gonorrhoeae*.

## Introduction

*Neisseria gonorrhoeae* (Gc), an obligate human pathogen and cause of the sexually transmitted infection gonorrhea, is an urgent public health threat that causes an estimated 86.9 million cases worldwide each year (1,2). Infection with Gc fails to elicit an effective host immune response, there is no protective immune memory to infection, a vaccine is not available, and resistance to antibiotics is increasing (3,4). There is only one remaining class of antibiotics that are recommended for treatment of gonorrhea, the cephalosporins, but strains resistant to ceftriaxone and cefixime have emerged, increasing the likelihood of untreatable gonorrhea (5),(6).

Gc infects human mucosal surfaces, including the female cervix and male urethra. At these sites, Gc stimulates a robust inflammatory response characterized by an abundant recruitment of neutrophils (7)(1). Although neutrophils produce and secrete antimicrobial components including proteases, cationic peptides, and reactive oxygen species, Gc is not cleared by the local neutrophil influx. Our group and others have identified several ways in which Gc resists killing by primary human neutrophils, including resistance to the antimicrobial proteins and reactive oxygen species that are made by activated neutrophils, limiting phagocytosis and phagosome maturation, and degradation of neutrophil extracellular traps (8). If neutrophilic inflammation is not resolved, it can cause irreversible tissue damage, leading to pelvic inflammatory disease and infertility (7).

Neutrophils associate with and internalize Gc through opsonic and non-opsonic means. The primary opsonins are antibodies and complement components, which are recognized by Fc receptors and complement receptors, respectively. Engagement of these receptors leads to internalization of the opsonized bacteria (9). The primary form of non-opsonic phagocytosis by neutrophils is through the interaction between the outer membrane opacity-associated (Opa) proteins and human carcinoembryonic antigen-related cell adhesion molecules (CEACAMs)(10,11). Isolates of Gc encode ≥ 9 distinct Opa proteins, and each *opa* gene is phase variable, generating extensive diversity in Opa expression within a bacterial population (12). Most Opa proteins bind one or more human CEACAMs. Expression of Opa proteins confers an advantage to the bacteria during cervical colonization by promoting interaction with the epithelial-expressed CEACAMs 1 and 5 (13). In contrast, Opa protein engagement of CEACAM-1 and CEACAM-3(10,14–17) on human neutrophils stimulates production of reactive oxygen species (ROS)(18), efficient bacterial binding and phagocytosis, and bacterial killing (19). Curiously, Opa-expressing (Opa+) Gc predominate in neutrophil-rich exudates from individuals with gonorrhea(20–22). It is an open question in the field why there is this discrepancy between *in vivo* and *ex vivo* observations with human neutrophils.

Inflammatory secretions are replete with human serum components as a result of serum transudate into the cervical lumen, which occurs under homeostatic conditions, and serum leakage due to breaching of the epithelial barrier, which occurs during the inflammatory conditions of Gc infection (7). One prominent serum component is the fluid-phase complement inhibitor C4b-binding protein (C4BP) (reviewed in (23)). The C4BP that primarily circulates in human blood is a 570 kDa multimer: 7 alpha (α) subunits (75 kDa, composed of 8 complement control protein (CCP) domains) and 1 beta (β) subunit (30 kDa, composed of 3 CCP domains). The C-terminal domains of the α and β chains mediate multimerization, and form a high affinity complex with the anticoagulation factor Protein S (PS) (7α1βPS). A 7α form of C4BP, which lacks a β chain and PS, is also present in the bloodstream, and increases in relative abundance compared to the 7α1βPS form of C4BP during conditions of inflammation. C4BP is present in human serum at an estimated concentration of 200 μg/mL(24). It inhibits complement activity in the classical and mannose-binding lectin pathways. CCP1 of the α chain binds complement component C4b to prevent assembly of the classical C3 convertase. C4BP also accelerates the decay of the C3 convertase by acting as a cofactor for Factor I, a serum protease that cleaves C3b and C4b.

C4BP binds to the surface of most strains of Gc and to other pathogenic bacteria including *Streptococcus pyogenes, Moraxella catarrhalis, Escherichia coli strain K1, Borrelia recurrentis, Haemophilus influenzae, Yersinia pestis, Bordetella pertussis*, and *Neisseria meningitidis*, through which it confers resistance to serum bactericidal activity in Gram-negatives and decreases complement-mediated phagocytosis in Gram-positives (25). The primary target of C4BP on Gc is the PorB porin; CCP1 of C4BP binds alleles of both PorB1a and PorB1b genotypes (26). Of 190 recently isolated clinical strains of Gc, 89.7% of PorB1a isolates and 19.3% of PorB1b isolates bound human C4BP(27). Commonly used lab strains FA1090 (genotype PorB1b) and 1291 (genotype PorB1b) bind C4BP(26). Gc does not bind C4BP of other species with the exception of chimpanzee, a species in which successful experimental gonorrhea infection has been accomplished, accounting in part for the human specificity of Gc infection(28). Roles of C4BP in the pathogenesis of Gc beyond inhibiting complement-mediated lysis have not been reported.

In this study, we made the unexpected observation that incubation of Opa+ Gc of strain FA1090 with normal human serum enhances its resistance to killing by primary human neutrophils and suppresses the neutrophil oxidative burst, independently of complement. We directly linked this phenomenon to a novel complement-independent function for C4BP. Using a derivative of FA1090 that constitutively expresses only the CEACAM-1 and -3 binding OpaD protein (OpaD+ Gc)(14), we found that binding of C4BP by Gc significantly enhances its survival from primary human neutrophils and suppresses neutrophil production of reactive oxygen species. Conversely, serum depleted of C4BP was unable to suppress the neutrophil oxidative burst and did not increase Gc survival after neutrophil challenge. C4BP addition reduced the association and phagocytosis of OpaD+ Gc by neutrophils. These outcomes all required binding of C4BP to the Gc surface, as shown using mutants of both C4BP and PorB1b that abrogate their interaction, and extended to variants of Gc expressing other Opa proteins. We propose that binding of C4BP enables Gc to avoid multiple facets of human innate immunity, by resisting neutrophil phagocytic killing as well as its canonical role in limiting complement-mediated lysis.

## Results

### Incubation of *Neisseria gonorrhoeae* with normal human serum limits neutrophil anti-gonococcal activity

To investigate the effect of serum on Gc-neutrophil interactions, OpaD+ Gc was incubated in pooled normal human serum, then exposed to adherent, IL-8 treated primary human neutrophils, and CFU were enumerated from cell lysates over time. We expected that serum would decrease survival of OpaD+ Gc by neutrophils via complement-mediated opsonophagocytosis. Instead, preincubation of the bacteria with serum significantly increased the number of OpaD+ Gc recovered from neutrophils, and in fact was equivalent to recovery of Opaless Gc at 30 minutes post-infection (**Fig. 1A**). The effect of serum on OpaD+ Gc was concentration-dependent, with OpaD+ Gc incubated in 10% and 25% serum surviving significantly better than bacteria without serum at 60 minutes (**Fig. 1B**). Serum had no significant effect on survival of OpaD+ Gc in the absence of neutrophils, reflecting the serum-resistant nature of the FA1090 genetic background (**Fig. S1)**.

**Figure 1.**
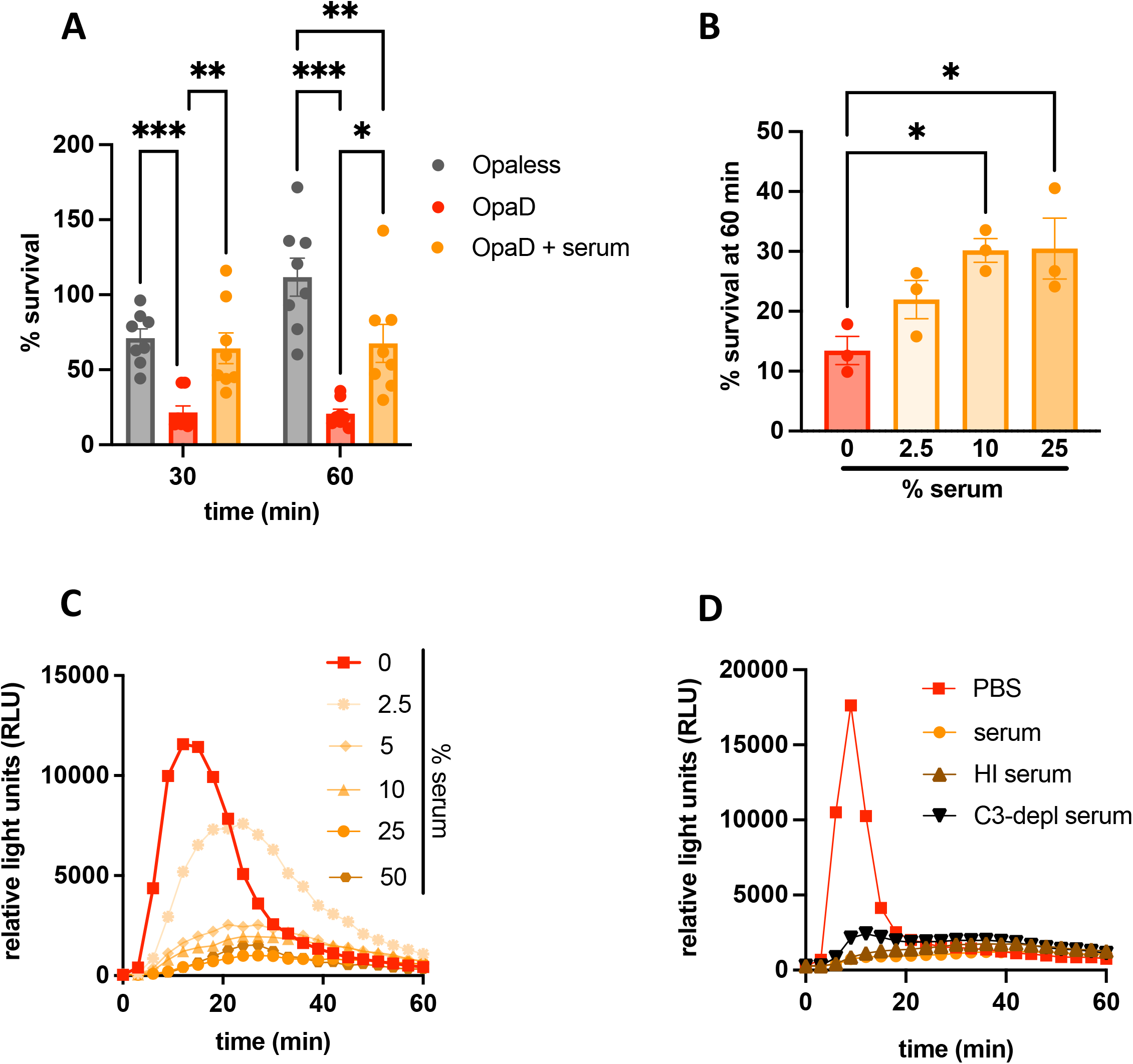
Incubation with normal human serum decreases neutrophil anti-gonococcal activity. (A) Adherent, IL-8-treated primary human neutrophils were exposed to Opaless Gc, OpaD+ Gc, or OpaD+ Gc that were preincubated in 25% normal human serum from healthy subjects. CFU were enumerated from neutrophil lysates at 0, 30, and 60 minutes (min) post infection, and bacterial survival is presented as the mean ± SEM of CFU at the indicated time point relative to the mean CFU for the same condition at 0 minutes for 8 independent experiments. Two-way ANOVA with Sidak’s post-hoc comparisons were used to compare each condition within each time point. In (B), OpaD+ Gc was mixed with the indicated percentage of normal human serum, and survival from neutrophils at 60 minutes was performed as in (A). One-way ANOVA followed by Tukey’s post-hoc comparisons was used to compare each condition to PBS alone (0% serum). *p<0.05, **p<0.01,***p<0.001. (C) OpaD+ Gc alone (PBS) or incubated in the indicated percentage of normal human serum was exposed to neutrophils in suspension at an MOI of 100 in the presence of luminol. ROS production was measured over the course of 60 minutes (min) as the relative light units (RLU) generated by luminol-dependent chemiluminescence. (D) ROS production was measured as in (C) from OpaD+ Gc incubated in serum, complement component 3-depleted serum (“C3-depl serum”), or heat-inactivated serum (“HI serum” (56 °C, 30 minutes)). (C-D) Results are one representative of 3 independent experiments.

We next examined how incubation with serum affects neutrophil functionality, using generation of reactive oxygen species (ROS) via luminol-dependent chemiluminescence as a readout. Preincubation of OpaD+ Gc with serum suppressed the resulting neutrophil ROS response in a concentration-dependent manner, with serum concentrations of 10% and higher abrogating ROS production (**Fig. 1C**). The suppressive effect of serum on neutrophil ROS production was replicated with a different isogenic derivative of Opaless Gc that constitutively expresses the Opa60 protein, which also binds to CEACAMs 1 and 3 (19,29), as well as Gc of strain 1291(26) expressing undefined Opa proteins (**Fig. S2**).

Given that complement is a major opsonic activity in serum, we tested whether the effects observed with serum on Opa+ Gc were complement-dependent. Heat-inactivated serum behaved identically to untreated serum in OpaD+ Gc-mediated suppression of neutrophil ROS (**Fig. 1D**) and retained the ability to enhance survival of OpaD+ Gc after exposure to neutrophils (see brown bars, **Fig. 4A**). Complement component C3-depleted human serum also suppressed the ability of OpaD+ Gc to elicit neutrophil ROS (**Fig. 1D**).

Together these results show that the presence of human serum unexpectedly increases the survival of Opa+ Gc from primary neutrophils and suppresses neutrophil activation, in a complement-independent manner.

### Identification of C4BP as the suppressive serum component

We characterized the component in serum that is responsible for suppressing neutrophil anti-gonococcal activity, using the reduction in ROS production as a readout. The suppressive activity was present in human serum but not from goat, mouse, rat, or cow (**Fig. 2A**). The suppressive component remained in the retentate of a 100 kDa molecular weight cutoff centrifugal filter device (**Fig. 2B**) and was sensitive to trypsinization (**Fig. 2C**), but the component was not immunoglobulin, as shown using IgG/IgA/IgM-depleted human serum **(Fig. 2D)**. Notably, the > 100 kDa retentate lost its ability to suppress ROS when it was preincubated with Gc, the bacteria pelleted, and the supernatant incubated with a new culture of OpaD+ Gc (“Gc-depleted” fraction) **(Fig. 2F)**. This observation enabled us to take an unbiased biochemical approach to identify the suppressive component from human serum, by following the reduction in neutrophil ROS production by OpaD+ Gc.

**Figure 2.**
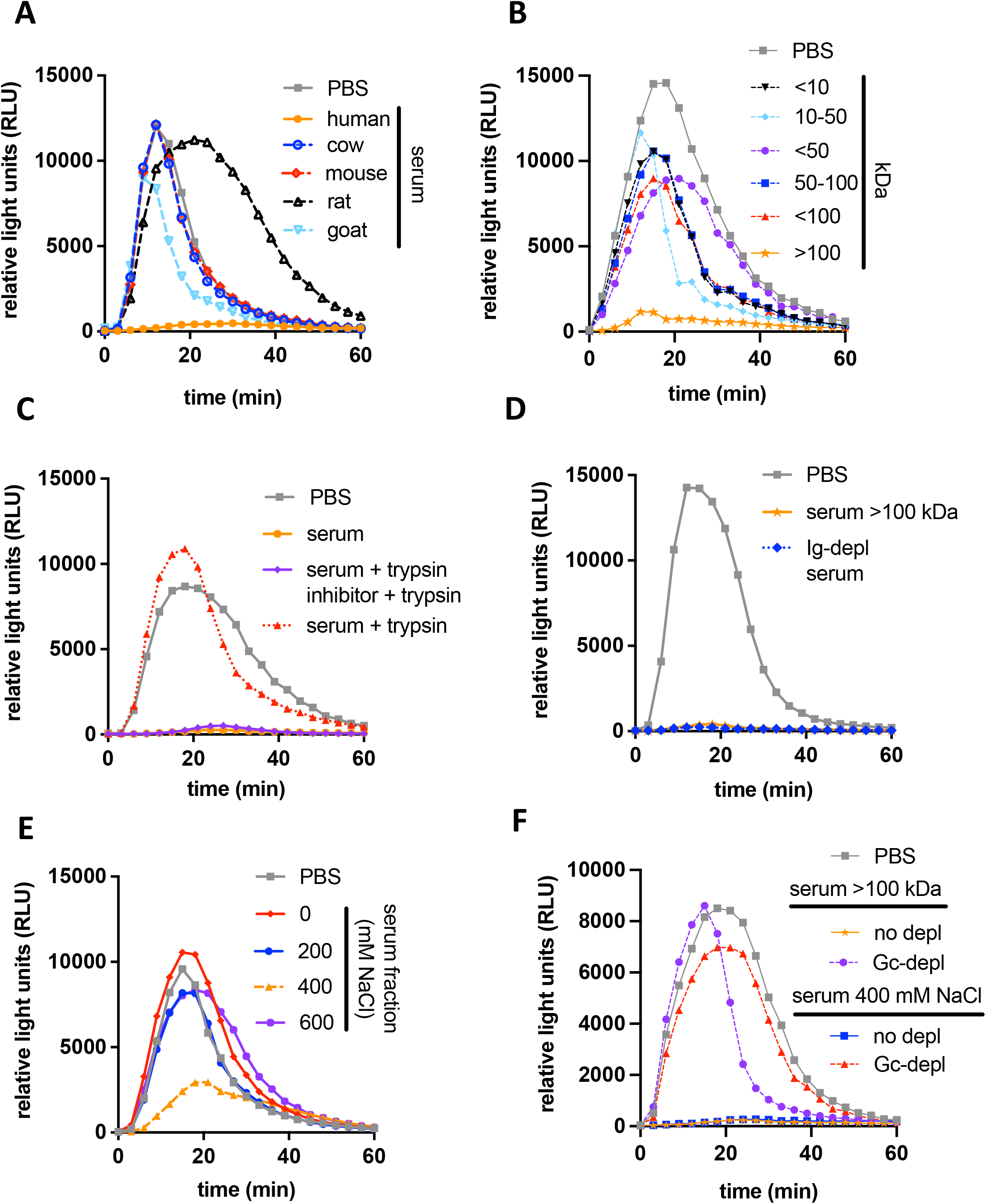
Characteristics of the serum component that suppresses Opa+ Gc induced neutrophil ROS. (A-F) OpaD+ Gc were exposed to primary human neutrophils and ROS production was measured as in Fig. 1C. Before addition to neutrophils, OpaD+ Gc was incubated with serum from the indicated species (A); the indicated molecular weight fraction of human serum (B); human serum that was intact (solid orange), treated with trypsin (dotted red), or treated sequentially with trypsin inhibitor then trypsin (purple) (C); the ≥ 100 kD human serum fraction from (B) (orange), or serum depleted of immunoglobulins G, M, and A (“Ig-depl serum”, dotted blue) (D); human serum fractions generated by anion exchange chromatography via elution with the indicated molarity of NaCl (E); or the indicated suppressive serum fraction from B and E that was pre-incubated with Gc (“Gc-depl”) or not (“no-depl”) (F). Incubation of OpaD+ Gc with PBS+ was used as a positive control for neutrophil ROS production in all conditions (grey).

To this end, pooled normal human serum was fractionated using anion exchange chromatography. Only the 400 mM NaCl eluate retained neutrophil suppressive activity (**Fig. 2E**) and was depletable by preincubation with Gc (**Fig. 2F**). The intact and Gc-depleted 400 mM fractions were trypsinized and analyzed by mass spectrometry (see **Methods**). The only peptide signatures that were present in abundance in the intact 400 mM fraction and absent from the Gc-depleted fraction corresponded to three proteins: C4BP α, C4BP β, and Protein S(30).

By imaging flow cytometry, C4BP was detectable on the surface of OpaD+ Gc that was incubated with intact or heat-inactivated human serum, or purified C4BP at 50 μg/mL (approximating the concentration of C4BP in 25% serum) (**Fig. 3A-C)**. No C4BP reactivity on OpaD+ Gc was detected using 25% serum that was C4BP-depleted (**Fig. 3B-C**). C4BP-depleted serum had heat-sensitive bactericidal activity against OpaD+ Gc, as expected for bacteria of strain FA1090(28) (**Fig. S1**). Going forward, C4BP-depleted serum and matched replete serum were heat-inactivated when mixed with Gc, such that any effects on Gc interactions with neutrophils were independent of complement-mediated lysis.

**Figure 3.**
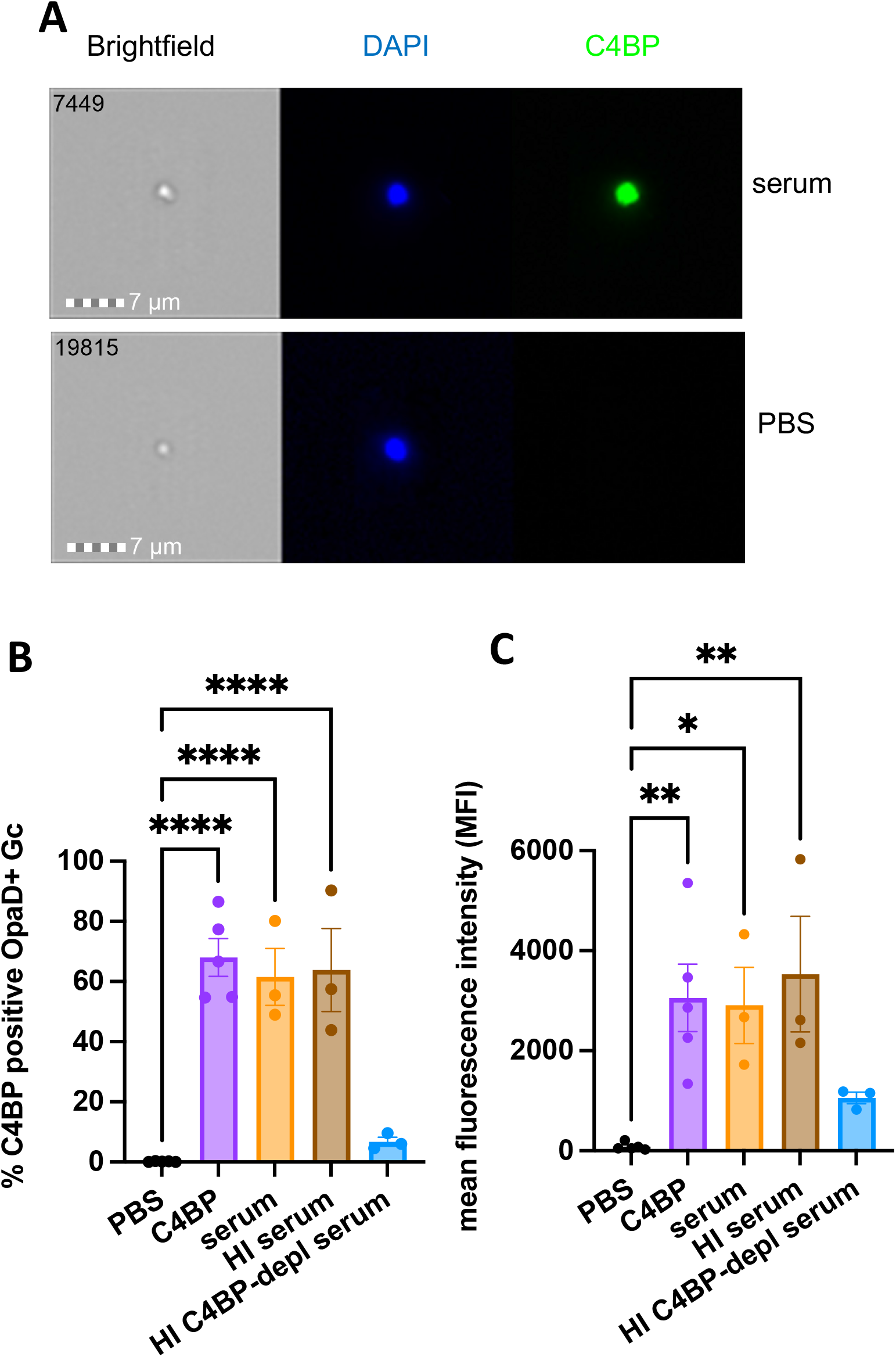
C4BP binds to OpaD+ Gc. OpaD+ Gc was incubated with 25% normal human serum (orange), heat-inactivated (HI) serum (brown), heat-inactivated C4BP-depleted (depl) serum (blue), or purified C4BP (50 μg/ml; purple) for 20 minutes at 37 °C. Gc was fixed and stained with DAPI for intact bacteria (blue) and for C4BP using anti-C4BP antibody followed by Alexa Fluor 488-coupled goat anti-rabbit IgG (green), and analyzed by imaging flow cytometry. (A) Representative images (image numbers 7449 and 19815) of a single serum-incubated (top) and PBS-incubated (bottom) bacterium. (B) is the percent C4BP-positive bacteria and (C) is the mean intensity of AF488 fluorescence of the total bacterial population. In B-C, results are the mean ± SEM of 3 independent experiments. Statistical analyses were performed using one-way ANOVA with Tukey’s post-hoc comparisons to PBS. *p<0.05, **p<0.01,****p<0.0001.

Together, these results led us to hypothesize that C4BP is the component of serum responsible for suppressing neutrophil anti-gonococcal activity and enhancing Gc survival from neutrophils.

### Binding of C4BP to *Neisseria gonorrhoeae* limits neutrophil anti-gonococcal activity

To test the contribution of C4BP in modulating Gc interactions with neutrophils, we made use of heat-inactivated C4BP-depleted serum and purified C4BP, each of which were compared with C4BP-replete serum. OpaD+ Gc that was incubated with C4BP-depleted serum survived just as poorly after exposure to human neutrophils as OpaD+ bacteria alone, and both survived significantly less well than bacteria incubated with C4BP-replete serum (**Fig. 4A**). Conversely, OpaD+ Gc that was incubated in purified C4BP survived similarly to Gc incubated with C4BP-replete serum, and both survived significantly better than untreated OpaD+ bacteria (**Fig. 4B**).

**Figure 4.**
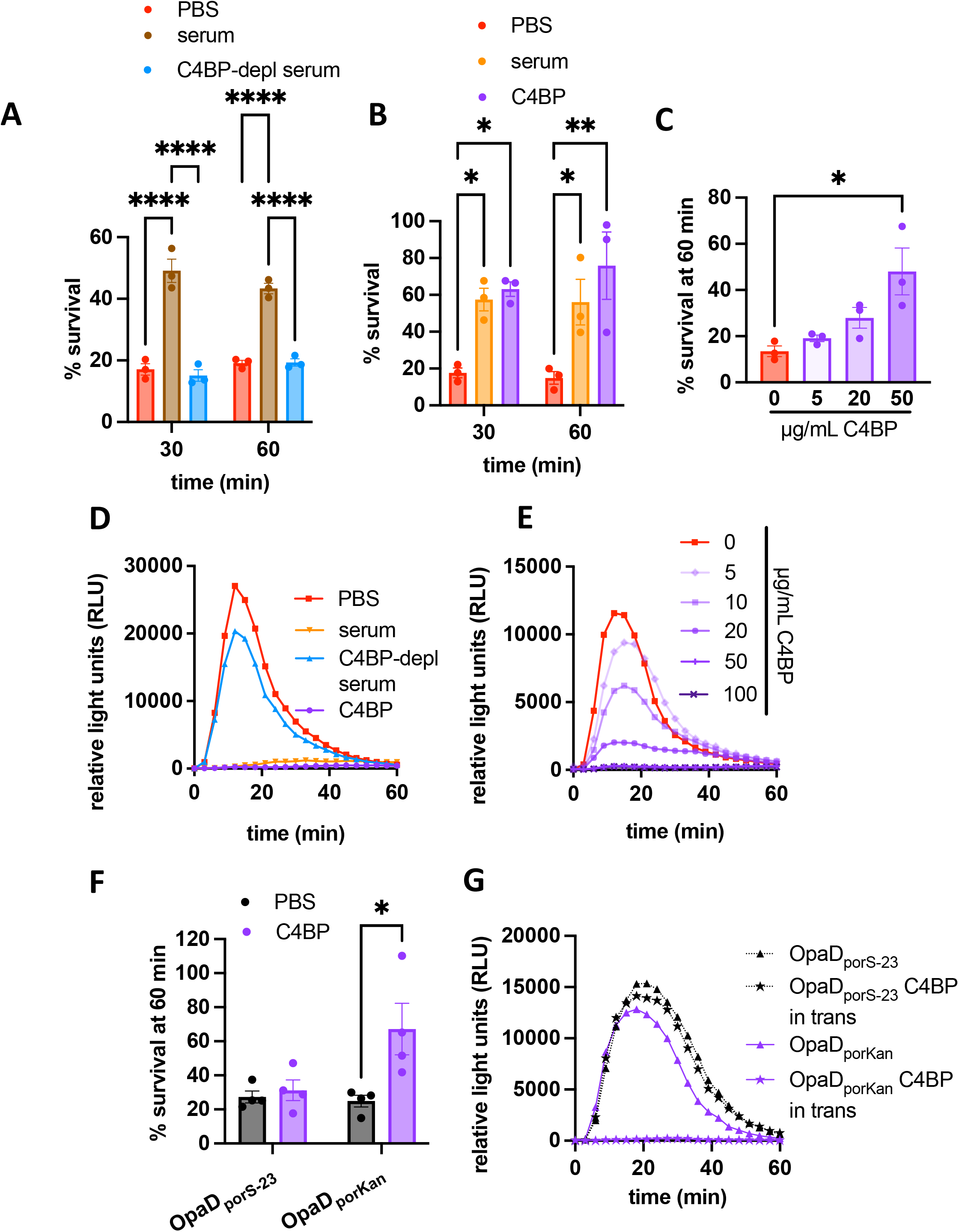
C4BP is necessary and sufficient for serum-mediated suppression of neutrophil anti-gonococcal activity. (A-C) Adherent, IL-8-treated neutrophils were exposed to OpaD+ Gc incubated in PBS+ (red) or the following (all at 25% final concentration) as in Fig. 1A: (A) heat inactivated serum that was C4BP-replete (brown) or C4BP-depleted (depl, blue); (B) C4BP-replete serum (orange) or 50 μg/mL purified C4BP (purple); (C) C4BP at the indicated concentrations. In (A-B), two-way ANOVA with Sidak’s post-hoc test was used to compare each condition within each time point for 3 independent experiments; (C) used one-way ANOVA followed by Tukey’s post-hoc comparisons to compare each condition to PBS alone (0 μg/mL C4BP). (D-E) Neutrophil ROS production was measured as in Fig. 1C in response to OpaD+ Gc that was incubated (D) in PBS, C4BP-replete serum (orange), heat inactivated C4BP-depleted serum (blue), or purified C4BP at 50 μg/mL (purple); or (E) in the indicated concentration of C4BP. (F) Neutrophils were exposed to OpaD_porS-23_ Gc or OpaD_porKan_ Gc, with or without C4BP addition to the infection milieu (50 μg/mL), and bacterial survival after 60 minutes was measured as in (A). Results are the mean ± SEM of 3 independent experiments; two-way ANOVA with Sidak’s post-hoc comparisons were used to compare each condition. *p<0.05, **p<0.01,****p<0.0001. (G) OpaD+ Gc (grey solid lines), OpaD_porS-23_ Gc (black dotted lines), and OpaD_porKan_ Gc (purple solid lines) alone (PBS; triangles), incubated with C4BP (open circles), or added to wells containing C4BP at a final concentration of 50 ug/mL (C4BP “in trans”; stars) were exposed to neutrophils, and ROS production was measured as in (E). Results in (D, E, and G) are one representative of 3 independent experiments.

As with normal human serum (**Fig. 1B**), incubation with purified C4BP enhanced the survival of OpaD+ Gc from neutrophils in a concentration-dependent manner (**Fig. 4C**). C4BP recapitulated the serum-mediated suppression of OpaD+ Gc-induced neutrophil ROS (**Fig. 4D**) in a concentration-dependent manner (**Fig. 4E**). Conversely, incubation of OpaD+ Gc in C4BP-depleted serum induced ROS release from neutrophils similarly to untreated OpaD+ Gc (**Fig. 4D**). As with OpaD+ Gc, incubation with C4BP significantly increased the survival of Opa60+ Gc from neutrophils (**Fig. S3A**), and neutrophil ROS elicited by Opa60+ Gc was suppressed by incubation with C4BP (**Fig. S3B**).

To investigate the structural requirements of C4BP for suppression of neutrophil anti-gonococcal activity, we utilized 3 different forms of C4BP (**S4A**): 7α1βPS (predominant form in plasma), 7α1β (no protein S), and 7α (plasma purified, no β chain or protein S). Each was recognized with anti-C4BPA antibody (**Fig. S4B**) and was able to bind to the surface of OpaD+ Gc (**Fig. S4C**). When mixed with OpaD+ Gc, all three forms of C4BP enhanced bacterial survival from neutrophils (**Fig. S4D**) and suppressed neutrophil ROS (**Fig. S4G)**, when compared with the effect of C4BP-depleted serum. Thus, the α chain multimer, which is made in inflammatory conditions, is sufficient to enhance OpaD+ Gc survival from neutrophils and suppress neutrophil ROS.

To directly test whether binding to Gc was necessary for the suppressive activity of C4BP on neutrophils, we took two complementary approaches. First, we used two forms of recombinant C4BP that have been reported, and which we validated, to be incapable of binding to Gc (**Fig. S4A**,**C**): 7α D15N/K24E, which contains two human-to-rhesus mutations in α chain CCP1 that abrogate PorB binding (31), and 1α, which is a monomer due to a deletion of the C-terminal linker and shows decreased binding to ligands due to loss of avidity (26). 7α D15N/K24E and 1α did not enhance OpaD+ Gc survival from neutrophils (**Fig. S4E-F)** or suppress neutrophil ROS release (**Fig. S4G**). Second, we engineered a strain of OpaD+ Gc with a mutant *porB* gene (see **Methods** for mutated residues) that was reported to not bind C4BP. We confirmed that the resulting mutant, OpaD_porS-23_, was unable to bind C4BP by imaging flow cytometry, while the isogenic control, OpaD_porKan_, bound C4BP similarly to OpaD+ Gc (**Fig. S5A-B**). OpaD_porS-23_ and OpaD_porKan_ Gc were incubated in medium containing purified multimeric C4BP, then added to neutrophils without washing away the C4BP (e.g. “in trans”).

Addition of C4BP in trans did not rescue the survival of OpaD_porS-23_ from neutrophils (**Fig. 4F**) and did not block the production of neutrophil ROS (**Fig. 4G**). However, C4BP added in trans did enhance the survival of OpaD_porKan_ Gc from neutrophils (**Fig. 4F**), and blunted the neutrophil oxidative burst that is stimulated by OpaD_porKan_ Gc similarly to OpaD+ Gc (**Fig. 4G)**. These results confirmed that the suppression of neutrophil antigonococcal activity by C4BP required its binding to Gc, and was not due to an unexpected, direct interaction of C4BP with neutrophils.

Taken together, these results show that C4BP is necessary and sufficient for serum-mediated enhanced survival of Opa+ Gc from neutrophils and for suppressing the oxidative burst that is elicited by Opa+ Gc, in a manner that is dependent on binding of C4BP to the bacterial surface.

### C4BP significantly decreases neutrophils’ association with and internalization of OpaD+ *Neisseria gonorrhoeae*

We recently reported that the main driver of Gc susceptibility to killing by neutrophils was the efficiency and extent of phagocytosis(19), in keeping with the finding that Gc internalized by neutrophils has significantly poorer survival than Gc that remains extracellular(15). Moreover, interaction of Gc with surface-expressed receptors such as CEACAMs can activate neutrophils, leading to release of ROS(18). For these reasons, we hypothesized that C4BP in serum enhances survival of Gc and suppresses ROS release from neutrophils by blocking bacterial binding and phagocytosis. To test this hypothesis, we measured the effect of C4BP on Gc association with and internalization by neutrophils over 1 hour by imaging flow cytometry (**Fig. 5A**)(32). In keeping with previous reports(14,15,19), OpaD expression promoted the rapid association and internalization of Gc with neutrophils (**Fig. 5B**,**C**), which was significantly reduced when OpaD+ Gc was mixed with normal human serum (**Fig. 5B-C**). This effect was C4BP-dependent, since OpaD+ Gc that was incubated in C4BP-depleted serum was indistinguishable from untreated bacteria in neutrophil association (**Fig. S6A**) and internalization (**Fig. 5D**), and both were significantly less than Gc incubated in C4BP-replete serum. Conversely, incubation of OpaD+ Gc with C4BP significantly decreased the percentage of neutrophils with associated (**Fig. S6B**) and internalized (**Fig. 5E**) OpaD+ Gc compared with untreated bacteria. The effect of serum and C4BP on internalization was predominantly driven by a reduction in bacterial interaction with the neutrophil surface, since incubation of Gc with serum or C4BP, compared with PBS as a control, did not affect the percentage of neutrophil-associated bacteria that were phagocytosed at 30 minutes (**Fig. 5F**). Gc binding to C4BP was required for these effects, since the addition of C4BP in trans had no effect on neutrophil association with (**Fig. S6C**) or internalization of (**Fig. 5G**) OpaD_porS-23_ Gc. Thus, C4BP is necessary and sufficient for the serum-mediated decrease in association and internalization of OpaD+ Gc by neutrophils.

**Figure 5.**
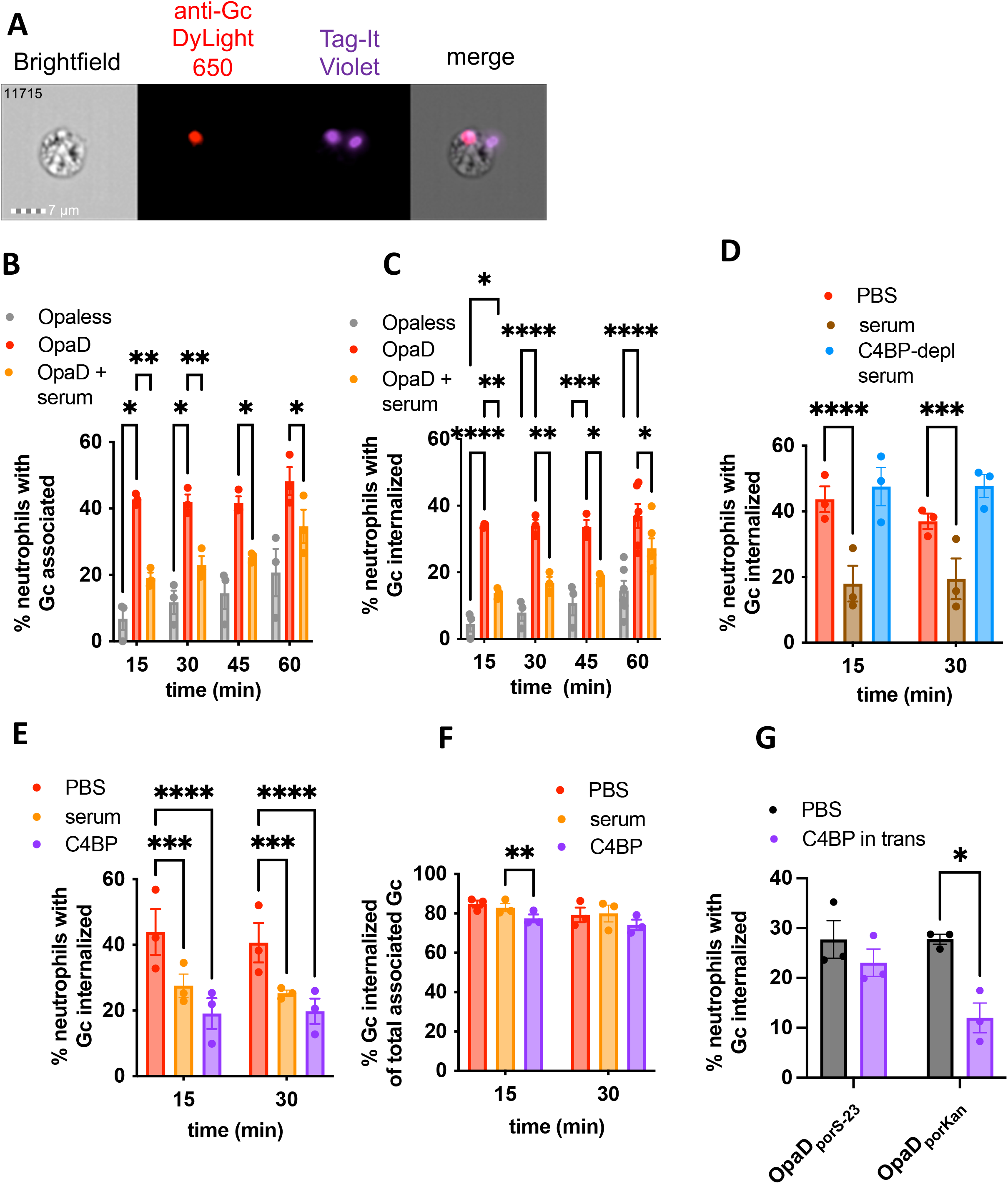
C4BP is necessary and sufficient for serum-mediated decrease in neutrophil internalization of OpaD+ Gc. Gc were labeled with Tag-IT Violet (TIV), treated as indicated in the following graphs, and incubated with adherent, IL-8-treated primary human neutrophils. At the indicated times, cells were fixed and stained with DyLight 650 (DL650)-labeled anti-Gc antibody without permeabilization to recognize extracellular bacteria. Neutrophils were analyzed via imaging flow cytometry. (A) Representative cell number 11715 of neutrophils with untreated OpaD+ Gc at 60 min, showing images captured from channels for (left to right) phase contrast, intracellular bacteria (red), total associated bacteria (purple), and merge. (B-C) Neutrophils were exposed to Opaless Gc (gray), OpaD+ Gc (red), or OpaD+ Gc incubated in C4BP-replete serum (25%, orange). (B) reports the percentage of single, focused, intact neutrophils with ≥1 cell-associated bacterium (TIV^+^); (C) reports the percentage of neutrophils with ≥1 phagocytosed bacterium (TIV^+^ DL650^−^). (D) Neutrophils were exposed to OpaD+ Gc in PBS (red), or heat-inactivated serum that was intact (brown) or C4BP-depleted (blue, both at 25% final), and the percentage of phagocytosed bacteria was calculated as in (C). (E-F) Neutrophils were exposed to OpaD+ Gc in PBS (red), serum (25%, orange), or purified C4BP (50 μg/ml, purple). In (E), the percentage of phagocytosed bacteria was calculated as in (C). In (F), the percentage of cell-associated Gc that are internalized was calculated. Results are the mean ± SEM from ≥3 independent experiments. Statistical analyses were performed by two-way ANOVA followed by Sidak’s multiple comparisons. (G) OpaD_porS-23_ Gc and OpaD_porKan_ Gc were incubated with neutrophils alone (grey bars) or with 50 μg/mL C4BP added to the media immediately prior to infection (“in trans”) for 30 minutes. The percentage of neutrophils with ≥ 1 phagocytosed bacterium was calculated as in (C) from 3 independent experiments. Statistical analyses were performed by two-way ANOVA followed by Sidak’s multiple comparisons. *p<0.05, **p<0.01,***p<0.001, ****p<0.0001.

**Figure 6.**
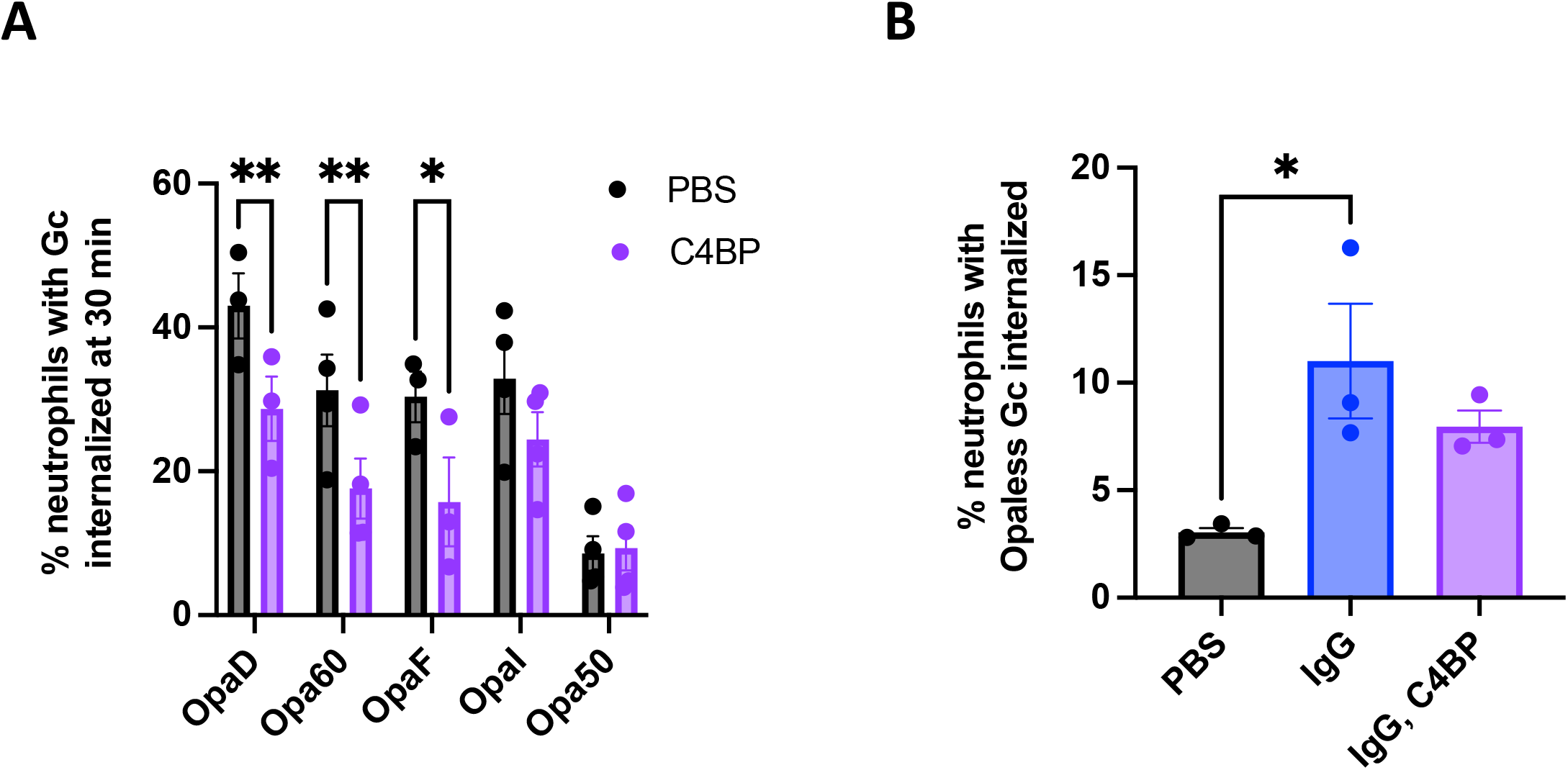
Binding of C4BP prevents internalization of CEACAM-binding Gc by neutrophils. (A) The indicated variants of Gc were incubated with C4BP (purple), or PBS (grey). The percentage of neutrophils with≥ 1 phagocytosed bacterium was calculated as in Fig. 5C. Data represent the mean ± SEM of ≥3 independent experiments. Statistical analyses were performed by two-way ANOVA followed by Sidak’s multiple comparisons. (B) Opaless Gc was incubated with C4BP (50 ug/mL), opsonized in rabbit anti-Gc IgG (80 μg/mL, 20 minutes, 37 °C), sequentially incubated with IgG then C4BP, or left untreated. Unopsonized Gc and IgG-opsonized Opaless Gc bound similar amounts of C4BP (**Fig. S8E**). The percentage of neutrophils with ≥ 1 phagocytosed bacterium was calculated as in (A). Statistical analyses were performed by one-way ANOVA followed by Tukey’s multiple comparisons. Opaless Gc incubated with C4BP behaved similarly to Opaless Gc (not shown). *p<0.05, **p<0.01.

To evaluate the effect of C4BP on the different routes of phagocytosis of Gc by neutrophils, we made use of a panel of single-Opa-expressing Gc that have different receptor-binding capacities and their Opaless parent (19). All isolates bound C4BP similarly (**Fig. S7A-D**). As we found for OpaD+ Gc, incubation of Opa60+ Gc (CEACAM-1 and CEACAM-3 binding) and OpaF+ Gc (CEACAM-1 binding) with C4BP significantly decreased bacterial association with and internalization by neutrophils (**Fig. S8A, 6A**). In contrast, C4BP binding to Opa50+ Gc (binds heparan sulfate proteoglycans (HSPGs) and not CEACAMs(16)) and OpaI+ Gc (CEACAMs 1 and 3 as well as HSPGs(19,33)) had no effect on interactions with neutrophils (**Fig. S8A, 6A**). The lack of effect of C4BP on Opa50+ Gc held true after 60 minutes of infection, where more neutrophils had associated and internalized the bacteria (**Fig. S8B-C**). Moreover, neutrophils’ association with and internalization of Opaless Gc, which does not express Opa proteins and does not bind CEACAMs or HSPGs, was similarly unaffected by incubation with C4BP at 60 minutes (**Fig. S8B-C**). To test if C4BP could block Gc-neutrophil interactions that are promoted by phagocytic receptors other than CEACAM family members, we used a Gc-specific IgG to drive phagocytosis of Opaless Gc through neutrophil Fc receptors. As expected, opsonization with IgG significantly increased the percentage of neutrophils with associated and internalized Opaless Gc, compared to unopsonized bacteria (blue vs. gray bars, **Fig. S8D, 6B**). However, adding C4BP had no effect on the ability of neutrophils to associate with or internalize IgG-opsonized Gc (purple bars, **Fig. S8D, 6B)**. Addition of IgG did not affect C4BP binding, and vice versa **(Fig. S8E)**. Together, these results suggest that the antimicrobial, antiphagocytic effect of C4BP on Gc is predominantly exerted on CEACAM-binding bacteria, and does not extend to Gc that are recognized by neutrophils by other means.

## Discussion

Our work uncovers an unexpected complement-independent role for C4BP in Gc pathogenesis: enhancing resistance to neutrophil clearance. Through an unbiased biochemical screen, C4BP multimer was identified as the complement-independent serum component that enhanced survival of Opa+ Gc from neutrophils and suppressed neutrophil activation, as read out by ROS production. The effects of C4BP were ascribed to inhibition of bacterial binding and phagocytosis by neutrophils. Use of both bacterial mutants and C4BP variants demonstrated that the effect of C4BP required binding to the bacterial surface. These effects were limited to Gc expressing Opa proteins that bind to some CEACAMs. Binding of C4BP may provide one explanation for why Opa+ Gc predominate in infected patient exudates, in spite of the potential for Opa proteins to activate neutrophils and promote bacterial killing *in vitro* in the absence of serum and C4BP. Our findings add novelty to an extensive literature demonstrating that many pathogens recruit C4BP to limit complement-mediated lysis, by showcasing that C4BP can also act in a complement-independent manner to thwart phagocytic killing **(Figure 7)**.

**Figure 7.**
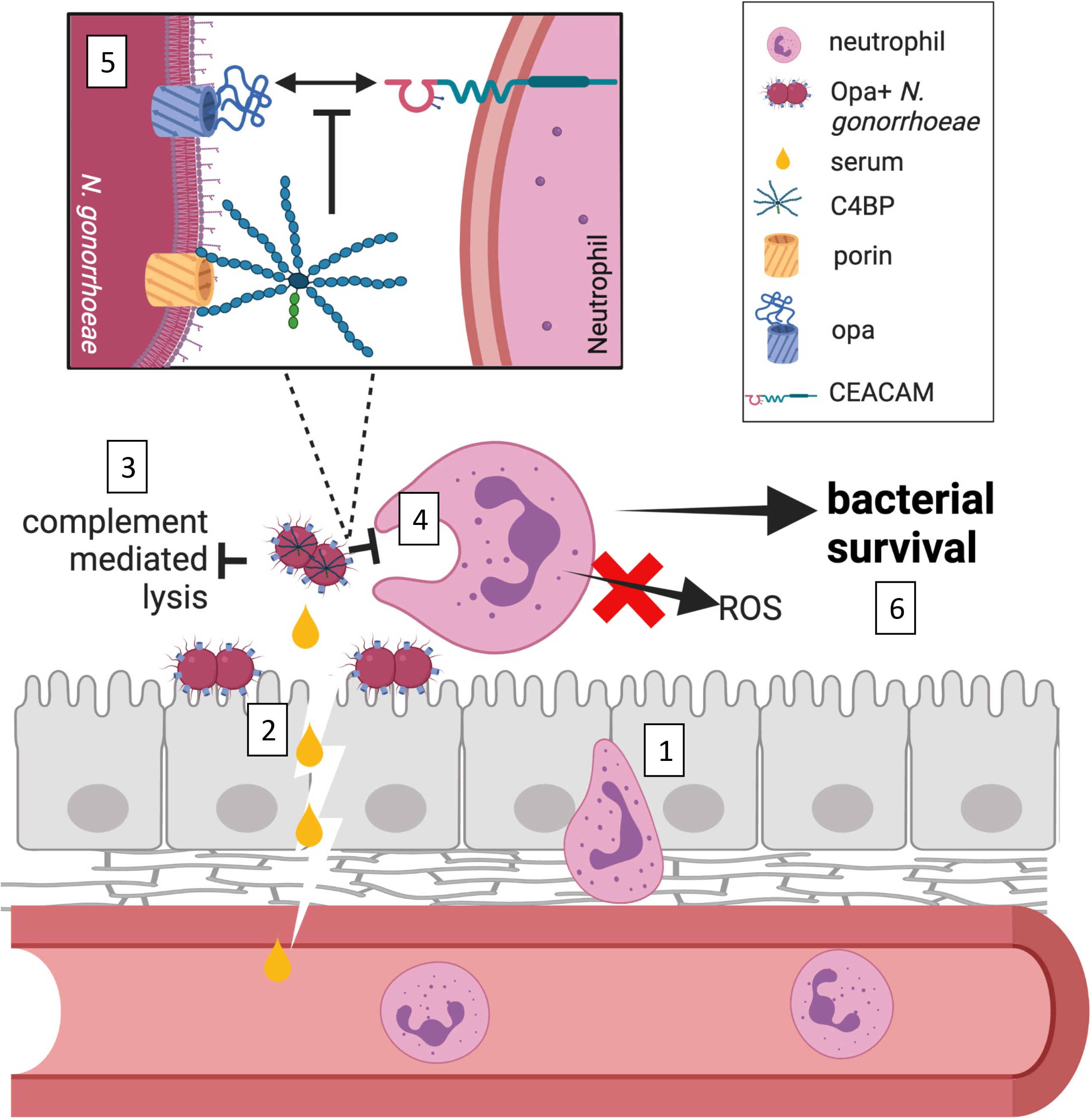
Complement-dependent and -independent roles of C4BP in gonococcal pathogenesis. Neutrophils are recruited to mucosal sites of *Neisseria gonorrhoeae* infection by extravasating from the blood stream and crossing the epithelia paracellularly [1]. Inflammatory conditions of the infection as well as serum transudate bring *N. gonorrhoeae* into contact with C4b-binding protein (C4BP) [2]. The ability of *N. gonorrhoeae* to bind to C4BP to inhibit complement-mediated lysis has been extensively documented [3]. This work shows that C4BP that is bound to the surface of *N. gonorrhoeae* decreases phagocytic uptake by neutrophils and suppresses neutrophil reactive oxygen species (ROS) production, in a complement-independent manner [4]. C4BP binding decreases the interaction of CEACAM-binding, opacity protein (Opa)-expressing *N. gonorrhoeae* with neutrophils, but not bacteria that are opsonized with IgG or that express Opa proteins that engage other ligands [5]. As a consequence, C4BP enhances survival of *N. gonorrhoeae* from neutrophils, a novel function for this canonical complement inhibitor [6].

Gc is expected to encounter C4BP in three conditions during infection. The first is during cervical infection, since serum transudate is present in the female genital tract. The C4BP concentration in the FRT is equivalent to approximately 11% complement, a value that is based on complement-based lytic activity of cervical fluid(34). We detected C4BP in human serum, and lower but measurable concentrations of C4BP in human mucosal fluids, including seminal plasma, vaginal fluid, menstrual blood, saliva, and tears (**Table S1**), implying Gc could be exposed to C4BP at the earliest stages of human infection at other sites as well. Even at C4BP levels well below what is found in serum, Gc would concentrate C4BP on its surface, as shown by the ability of the bacteria to deplete C4BP from fractionated human serum. The second situation where Gc would encounter C4BP is in conditions of inflammation in acute gonorrhea, where serum leakage occurs due to mucosal damage and breaches from the influx of neutrophils. Inflammatory conditions also increase circulating levels of the 7α form of C4BP(35),(36).Interestingly, C4BP 7α has been found reduce innate immune inflammation in the context of systemic lupus erythematosus(37), another non-canonical role for C4BP. If 7α has similar anti-inflammatory properties in gonorrhea, future studies could investigate how different forms of C4BP are generated and function in the context of human disease. In the third scenario, Gc would encounter C4BP in the bloodstream during a disseminated gonococcal infection. Future studies could utilize transgenic animal models to demonstrate the effect of human C4BP on infection outcome in each of these scenarios.

The ability of Gc to survive exposure to neutrophils correlates with its resistance to phagocytosis(19). We found here that incubation with C4BP reduces neutrophil association with and internalization of Opa+ Gc, which can explain how C4BP limits killing of Gc by neutrophils in this infection model. However, C4BP may also enhance Gc resistance to neutrophils by inhibiting neutrophil signaling pathways that lead to release of antimicrobial products. This is in line with our finding that C4BP incubation also suppressed the ability of Opa+ Gc to elicit neutrophil ROS, even though ROS does not contribute to neutrophil anti-gonococcal activity(15,38). Neutrophil NADPH oxidase activation requires a series of cytoplasmic phosphorylation events, Rac GTPase activation, and granule release, all of which contribute to the overall activation state of neutrophils (18). Moreover, release of anti-gonococcal proteases from neutrophil primary granules requires signaling via nonreceptor tyrosine kinases like Src and Syk(39,40). Effects of C4BP on these pathways would be expected to work in concert with C4BP’s antiphagocytic effect to enhance Gc survival from neutrophils.

Our results suggest that C4BP enhances Gc resistance to neutrophil killing by sterically hindering bacterial binding and phagocytosis by neutrophils. Curiously, this was limited to Opa-CEACAM interactions on the neutrophils, and did not affect phagocytosis of HSPG-binding Opa+ Gc or IgG-opsonized Gc. We note that PorB, the primary target for C4BP on Gc, is not known to be a ligand for any neutrophil receptors. Our data suggests that C4BP is not interfering with Opa-CEACAM binding by occluding the binding interface, as shown with a bacterial N-CEACAM precipitation assay **(Fig. S9A-B)**. Alternatively, C4BP could bind to a target on the neutrophil to send signals that prevent phagocytosis, but such a target has not been described. While protein S can bind to phosphatidylserine residues on apoptotic cells(41), multimeric C4BP lacking protein S still inhibited Gc phagocytosis and ROS production, suggesting this mode of interaction is not at play. Also, addition of C4BP to the porB_S-23_ non-binding mutant Gc did not suppress neutrophil functions, suggesting that soluble C4BP does not directly engage neutrophils to account for these effects.

It is noteworthy that serum incubation had the most pronounced effect on Gc-neutrophil association at early time points and was minimized over time. There are two nonexclusive explanations for this observation. One, if C4BP effects on nonopsonic phagocytosis are predominantly by steric hindrance, interaction between Gc and neutrophils could be expected to “catch up” over time. Two, release of neutrophil proteases could degrade C4BP on the surface of Gc. This could occur for Gc bound to neutrophils, if the proteases are released extracellularly, or phagocytosed bacteria, if released into the phagosome. Interestingly, an intracellular role for C3 in activating antibacterial autophagy for cytoinvasive bacteria has been described(42). It is intriguing to consider the possibility that C4BP on the surface of phagocytosed Gc could help protect the bacteria from intraphagosomal killing, potentially helping to explain how viable, intact Opa+ Gc are found inside neutrophils recovered from infected individuals (43,44).

This work has implications for the development of new antimicrobials and vaccines for gonorrhea, which are urgently needed for global public health. The Ram group has recently developed therapeutics for Gc that make use of bacterial-binding C4BP CCPs that are fused to IgG or IgM, which recruit complement components to the bacterial surface and elicit serum bactericidal activity(27). It is curious to consider how the C4BP effects on neutrophils reported here would affect the function of these therapeutics as treatments for gonorrhea. While IgG-fused C4BP might facilitate bacterial phagocytosis, the multimeric IgM fusion might block interactions with neutrophils rather than stimulate it, even if there is complement deposition.

Moreover, in light of efforts to develop a gonococcal vaccine, it would be crucial that a vaccine candidate can still bind Gc and recruit complement components in the presence of C4BP. If opsonophagocytosis contributes to vaccine-mediated protection, a vaccine candidate would need to overcome not only the effect of C4BP on limiting complement-dependent phagocytosis, but also the effect of C4BP on nonopsonic Opa-driven interactions between Gc and neutrophils. Since many gonorrhea vaccine studies use mouse sera, and mouse C4BP does not bind to Gc, these results highlight the need to address how C4BP, as well as other complement-limiting factors like factor H and lipooligosaccharide sialylation, would affect the efficacy of a vaccine candidate.

To our knowledge, C4BP has never been reported to enhance the survival of pathogenic bacteria in a complement-independent manner. In fact, the only report to date of C4BP affecting host-pathogen interactions in a complement-independent manner is a report from Varghese and colleagues showing that C4BP restricted viral entry of Influenza A subtype H1N1 into lung epithelial cells(45). It is intriguing to consider that the many pathogenic bacteria that recruit C4BP to their surface may do so to prevent complement-dependent and -independent neutrophil responses. This may be especially important for pathogenic bacteria like Group A streptococci that, like the pathogenic *Neisseria*, stimulate a potent pyogenic response(46). The protection afforded to Opa+ Gc by C4BP binding could allow Gc to express Opa proteins for the benefit of tight adherence to epithelial cells during infection, while decreasing the Opa-induced activation of neutrophils that would occur during the peak of inflammation. The complement-independent role for C4BP reported here shifts our understanding of the relationship between the cellular and noncellular components of the innate immune system, and how pathogens like Gc can exploit these factors to evade immune clearance for successful infection.

## Methods

### Biological fluids and proteins

#### Normal human serum preparation

Venous blood from 10 healthy human subjects, who were consented under a protocol approved by the University of Virginia Institutional Review Board for Health Sciences Research (IRB-HSR #13909), was collected into serum vacutainer blood collection tubes (BD 366430). After incubation at 37°C for 30 minutes, the serum supernatant was collected, pooled, diluted to 50% with PBS with 100 mg/L CaCl_2_ and 100 mg/L MgCl_2_ (“PBS+”; Gibco 14040133), and passed through a 0.2 μm filter. Serum was aliquoted and stored at -20°C and thawed on ice.

#### C4BP-depleted serum and matched C4BP-replete serum

C4BP was depleted from pooled healthy human serum as previously described(47) by passing fresh serum through a column coupled with MK104, a mouse mAb directed against CCP1 of the α-chain of C4BP, with C1q addback (80 μg/mL). Serum was stored at -80°C and thawed on ice. C4BP-replete serum (untreated serum from the same donor pool) was used for comparison. Serum was collected with written consent under ethical permit from Lund University (2017-582). C4BP depletion was verified via western blot and mass spectrometry (**Table S1**).

#### Modified sera

IgG/M/A-depleted serum (Sigma-Aldrich) was reconstituted from a lyophilized powder and diluted to 25% in PBS+. Complement C3-depleted serum was sourced from Complement Technologies (A314).

To create fractions of serum, pooled normal human serum was separated using size-exclusion centrifugal filter units (Amicon). For the <10 kDa fraction, the flow-through of a 10 kDa molecular weight cut off (MWCO) filter was collected. For the >100 kDa fraction, the retentate of a 100 kDa MWCO filter was collected and brought up to original volume with PBS+, and the flow-through collected as the <100 kDa fraction. The <50 kDa fraction was collected from the flow-through of a 50 kDa MWCO filter. For the 10-50 kDa fraction, the retentate of a 10 kDa MWCO filter was subsequently added to a 50 kDa MWCO filter, and the flow-through was collected. For the 50-100 kDa fraction, the retentate of a 50 kDa MWCO column was subsequently added to a 100 kDa MWCO filter, and the flow-through was collected. Serum fractions were sterilized by passage through a 0.2 μM filter and stored at -20° C.

For digestion, pooled normal human serum, diluted to 25% in PBS+, was incubated with 1.5 mg/mL trypsin (Sigma T6763) for 2 hours at 37 °C. Serum was placed on ice, soybean trypsin inhibitor (3 mg/mL) (Gibco 17075-029) was added, and the serum was used immediately after preparation.

#### Other animal sera

Normal rat serum (Enco Scientific Services 13551), fetal bovine serum (HyClone SH30071.03, Lot AC10223670), and normal goat serum (Gibco, 16210-064) were commercially sourced. Fresh normal mouse serum was a gift from the lab of Jonathan Kipnis (formerly of UVA).

#### Non-serum human biological fluids

The following human biological fluids were sourced from Lee Biosolutions, Inc: Vaginal fluid (991-10-P-1), seminal plasma (991-04-SPP-1), menstrual blood (991-15-P), tears (991-12-P), and saliva (991-05-P-PreC). All were stored at -20°C, thawed on ice, and centrifuged to remove insoluble material prior to use.

#### Purified and recombinant C4BP

Human serum-purified C4BP was sourced from Complement Technologies, Inc (A109). It was stored at -80°C, thawed on ice, and diluted in PBS+ to the desired concentration for experimentation.

C4BP 7α1βPS that was used in experiments with C4BP-depleted and -replete human serum was purified from human plasma by affinity chromatography using a MK104 antibody column as previously described(48). C4BP 7α1β was purified from plasma via a standard method that uses barium chloride precipitation followed by anion exchange chromatography and gel filtration. Protein S was then removed after incubation with ethylene glycol, which breaks the hydrophobic bond between C4BP and protein S, followed by affinity chromatography on heparin-Sepharose (49),(50). C4BP 7α, the species lacking the βchain, was purified from the supernatant of the barium chloride precipitation(49) using affinity chromatography with MK104. Recombinant forms of C4BP were expressed in human embryonic kidney (HEK) 293 cells. Recombinant multimeric C4BP 7α was purified via MK104 column as previously described(31). Recombinant C4BP 1α, the monomeric form of C4BP, has a deletion of the C-terminal linker that is required for multimerization, and was purified as previously described(26). Recombinant C4BP 7α D15N/K24E, in which CCP1 has two human-to-rhesus mutations in CCP1 that abrogate binding to Gc, was purified as previously described(31).

### Bacterial strains

This study used piliated FA1090 Gc that is deleted for all *opa* genes (Opaless(14)) and containing constitutively expressed versions of OpaD(14), Opa60(51), OpaI(19), or Opa50(51), which have a mutated signal sequence that does not phase-vary. Predominantly OpaF+ Gc was selected phenotypically based on colony opacity from strain *ΔopaBEGK* (14). Expression of OpaF was verified by Western blot using an OpaF-specific antibody (a gift from Marcia Hobbs, UNC)(52). Opa+ Gc from strain background 1291 was selected visually by colony opacity.

For all assays, Gc was grown overnight on gonococcal medium base (GCB, Difco) with Kellogg’s supplements I and II(53) at 37°C with 5% CO_2_. Piliated colonies were selected the next morning and grown for 8 hours on GCB, then grown in gonococcal base medium liquid (GCBL) culture with Kellogg’s supplements I and II overnight with rotation with two back dilutions the next day as described previously(54). Piliated Gc were enriched by using sedimented bacteria for the final back dilution.

OpaD_porS-23_ was generated by transforming OpaD+ Gc with genomic DNA from FA1090 S-23 (from Hank Seifert, Northwestern University). FA1090 S-23 contains 5 residues (amino acids 254-259) mutated to alanine in loop 6 of *porB* (Y254A, G255A, M257A, S258A, G259A), with a kanamycin resistance cassette inserted downstream(55). Transformants were selected on GCB containing 50 μg/mL kanamycin. *porB* was amplified from transformants by PCR using PorI (GGCGAATTCCGGCCTGCTTAAATTTCTTA) and PorII (GCGAAGCTTATTAGAATTTGTGGCGCAG), using conditions described in (56), and sent for commercial Sanger sequencing using the same primers. Isolates with the mutated *porB* were verified to not bind C4BP by flow cytometry (see <u>Detection of C4BP on bacteria</u>). OpaD_porKan_ is isogenic to OpaD_porS-23_ but retains the parental FA1090 *porB*.

### Neutrophil purification

Neutrophils were purified from venous blood of healthy human subjects who signed informed consent in accordance with UVA IRB-HSR protocol #13909, using dextran sedimentation followed by a Ficoll (Cytiva) gradient and erythrocyte lysis, as described previously(54,57). Neutrophils were resuspended in 1x PBS (no calcium or magnesium) (Gibco 14190-144) containing 0.1% glucose, stored on ice, and used within 2 hours of purification.

### Serum/C4BP incubation

Serum or C4BP was diluted in PBS+ to the desired final concentration. Gc was suspended in serum or C4BP, at a final volume of 250 μL for 10^8 Gc, for 20 minutes at 37°C. Bacteria were pelleted, the supernatant was discarded, and the pellet was washed in PBS+. No free serum or C4BP was present in experiments unless otherwise noted.

### Serum bactericidal activity assay

OpaD+ Gc (10^8) was mixed (see <u>Serum/C4BP incubation</u>) for 20 minutes in the indicated percentage of serum or C4BP-depleted serum, with or without heat inactivation at 56°C for 30 minutes. Bacteria were pelleted, washed, diluted, and plated on GCB agar. Colony forming units (CFU) were enumerated after overnight growth.

### Mass spectrometry

#### Identification of C4BP

Pooled normal human serum (see above) was fractionated using strong anion exchange chromatography with a HiTrap Q HP column (Cytiva). Fractions eluted with increasing concentrations of salt were tested for ROS suppressive activity (see <u>Neutrophil Assays</u> below). The suppressive fraction (400 mM NaCl) and activity-depleted fraction (400 mM NaCl preincubated with 5×10^8 OpaD+ Gc for 30 minutes at 37°C) were analyzed by mass spectrometry (MS) and tandem mass spectrometry (MS/MS) at the Biomolecular Analysis Core Facility at the University of Virginia. Samples were reduced with 10 mM DTT in 0.1 M ammonium bicarbonate followed by alkylation with 50 mM iodoacetamide in 0.1 M ammonium bicarbonate (both room temperature for 0.5 h). The samples were then digested overnight at 37°C with 0.1 μg trypsin in 50 mM ammonium bicarbonate. The samples were acidified with acetic acid to stop digestion and then purified using magnetic beads. The solution was evaporated for MS analysis.

The LC-MS system consisted of a Thermo Q Exactive HF mass spectrometer system with an Easy Spray ion source connected to a Thermo 3 μm C18 Easy Spray column (through pre-column). The extract (1 μg) was injected and the peptides eluted from the column by an acetonitrile/0.1 M acetic acid gradient at a flow rate of 0.3 μL/minute over 1.0 hours. The nanospray ion source was operated at 1.9 kV. The digest was analyzed using the rapid switching capability of the instrument acquiring a full scan mass spectrum to determine peptide molecular weights followed by product ion spectra (10 HCD) to determine amino acid sequence in sequential scans. This mode of analysis produces approximately 25000 MS/MS spectra of ions ranging in abundance over several orders of magnitude.

The data were analyzed by database searching using the Sequest search algorithm(58) against Uniprot Human. Scaffold Viewer software (Proteome Software, Inc) was used to visualize and compare the relative abundance of peptides in each fraction.

#### Detection of C4BP in secretions

We performed an untargeted data-dependent analysis of tryptic peptides of purified C4BP to identify target peptides and transitions for parallel reaction monitoring (PRM) analysis along with literature and PeptideAtlas (http://www.peptideatlas.org/) searches. Peptide LNNGEITQHR from C4BPα was selected, and a C-terminal isotope-labeled (“heavy” peptide) version was synthesized (Thermo Fisher Scientific).

Samples from distinct fluids were prepared for mass spectrometry analysis as follows. Proteins from 100 μL of each sample were precipitated by adding 1 mL of cold Methanol/Acetone (9:1) and incubated at -80 °C. Samples were centrifuged and washed twice with 1 mL of cold methanol. Protein pellets were reconstituted in 100 mM ammonium bicarbonate and total protein was measured using BCA. 10 μg of total protein of each sample were reduced with 10 mM DTT followed by alkylation with 50 mM IAA both at room temperature for 30 min. Trypsin digestion was performed using 1:25 enzyme to protein ration at 37 C for 16 h. Samples were acidified using acetic acid, peptides were purified using C-18 ZipTips and dried under vacuum. Each sample was reconstituted in 10 μL of formic acid containing 1 fmol/μL of the isotope-labeled peptide.

Peptide mixtures were analyzed on Thermo Orbitrap Exploris 480 system coupled to an EASY-nLC 1200 system. 1 μL of each sample was automatically injected into a Thermo 3 μm C18 Easy Spray column (through pre-column) and peptides eluted from the column by acetonitrile/0.1 % formic acid gradient at a flow rate of 300 μL/min over 68 minutes. The nanospray ion source was operated at 1.5 kV and the mass spectrometer was operated on positive mode acquiring targeted MS2 scans using Orbitrap resolution at 60,000, AGC target 300, isolation window of 1.2 m/z, and optimized collision energy for HCD of 20.

For quantification, the total peak areas of each peptide were used to obtain light to heavy ratio (L:H). Since the spiked-in heavy concentration is known, the L:H ratios were used to calculate the concentration of the peptide corresponding to the endogenous C4BP in each sample.

### Western Blotting

Proteins or Gc were boiled for 5 minutes in reducing sample buffer containing 60 mM Tris pH 6.8, 25% glycerol, 0.4% SDS, 5% β-mercaptoethanol, and 0.1% bromophenol blue. Proteins or lysates were resolved on a 4-20% gradient SDS-polyacrylamide gel (Criterion TGX 5671094, Bio-Rad), transferred in Towbin(59) buffer to nitrocellulose membrane, and the membrane was incubated in 5% BSA in Tris buffered saline with 0.1% Tween for 1 hour (blocking buffer). C4BP was detected using rabbit anti-C4BPA antibody (Novus Biologicals 88262) followed by goat anti-Rabbit H+L Alexa Fluor 680 cross adsorbed secondary antibody (Invitrogen A21109), and Opa or porin (loading controls) were detected with mouse anti-Opa 4B12 and mouse anti-porin H5.2 respectively followed by goat anti-Mouse IgG (H+L) Dylight 800 cross adsorbed secondary antibody (Invitrogen SA5-10176). All antibodies were diluted in blocking buffer. Bands were visualized via LI-COR Odyssey.

### Detection of C4BP on bacteria

Gc (10^8) was pelleted, washed, and incubated with 50 μg/mL C4BP diluted in PBS+ for 20 minutes at 37°C. Gc were pelleted, washed, and resuspended in 5 μg/mL rabbit anti-C4BPA antibody (Novus Biologicals 88262), diluted in RPMI + 10% FBS, for 30 minutes at 37°C. Gc were pelleted and resuspended in 5 μg/mL goat anti-mouse AF488 (Invitrogen, Carlsbad, CA) diluted in RPMI + 10% FBS for 30 minutes at 37°C. The samples were fixed in 2% PFA with 5 μg/mL 4′,6-diamidino-2-phenylindole (DAPI) for visualization of the bacteria. Samples were analyzed by imaging flow cytometry using Imagestream^X^ Mk II with INSPIRE^®^ software (Luminex Corporation) at the Flow Cytometry Core Facility at the University of Virginia, and data were analyzed using IDEAS® software.

### N-CEACAM binding to Gc

Binding of recombinant N-termini of CEACAM1 or CEACAM3 to Gc, with and without incubation with C4BP, was measured by imaging flow cytometry, based on fluorescence of mouse anti-GST antibody p1A12 (Biolegend) as in(29).

### Neutrophil Assays

#### Gc survival from neutrophils

Gc survival from primary human neutrophils was measured as described previously(54). In brief, 10^6 adherent, IL-8 treated neutrophils were synchronously exposed to 10^6 Gc (MOI 1) by centrifugation in RPMI containing 10% fetal bovine serum. After incubation at 37°C with 5% CO_2_ for the indicated times, neutrophils were lysed with 1% saponin and lysates were serially diluted and plated on GCB. Colony forming units (CFU) were enumerated after overnight growth, and survival was calculated as a percentage relative to the CFU enumerated at the 0 minute time point.

#### ROS measurement

ROS release from neutrophils was measured as described previously(18) by suspending neutrophils in Morse’s Defined Medium (MDM)(60) (1) with luminol in a white-bottomed 96 well plate (Falcon 353296). Bacteria were added to neutrophils at an MOI of 100. Luminescence was measured every 3 minutes over 1 hour using a VICTOR3 Wallac luminometer (Perkin-Elmer).

#### Neutrophil association with and internalization of Gc by imaging flow cytometry

Following the protocol outlined in (32), 2×10^6 adherent, IL-8 treated neutrophils were synchronously infected with 2×10^6 Tag-It Violet stained Gc by centrifugation (MOI 1). After incubation for the indicated times at 37°C with 5% CO_2_, paraformaldehyde was added (final concentration 2%) and cells were lifted with a cell scraper (Corning 353085). Cells were washed in PBS and blocked in 10% normal goat serum. Extracellular associated bacteria identified by reaction with rabbit anti-Gc antibody (Meridian B65111R) that was conjugated in-house with 1 μg/mL Dylight 650 (Thermo Scientific) according to manufacturer’s recommendations. Results are expressed as the percentage of neutrophils with at least 1 associated (bound and/or internalized; Tag-IT Violet+ Dylight 650+) Gc and the percentage of neutrophils with at least 1 internalized (Tag-IT Violet+ Dylight 650-) Gc.

### Statistics

Survival, binding, association, and internalization results are displayed as the mean ± SEM for n≥ 3 biological replicates, conducted on different days with different bacterial cultures and subjects’ neutrophils. Statistical comparisons were performed using one-way ANOVA followed by Tukey’s multiple comparisons post-hoc test, two-way ANOVA followed by Sidak’s multiple comparisons post-hoc test, or paired Student’s *t*-test as appropriate, using Prism GraphPad. For data sets analyzed by *t-*test or two-way ANOVA, data from the same biological replicate are paired, to account for inter-subject neutrophil variation. ROS results are presented as one representative graph of n≥3 biological replicates, which cannot be averaged because of day-to-day differences in the magnitude of luminescence emitted.

## Acknowledgements

This work was supported by NIH R01AI097312 and R21 AI157539 to AKC. LMW was supported in part by the UVA Wagner Fellowship and NIH T32 AI007046. AMB was supported by Swedish Research Council (2018-02392). MBD was supported by NIH Ruth L. Kirschstein Postdoctoral Fellowship (F32GM136076). JCR was supported in part by NIH T32 AI055432 and NIH T32 GM007267. We thank Louise Ball, Samuel Clark, and Christopher Baiocco, who performed preliminary experiments that informed this project. We thank Mike Solga of the UVA Flow Cytometry Core Facility (RRID: SCR_017829) and Nick Sherman, PhD of the UVA Biomolecular Analysis Facility for their advice. We thank Hank Seifert, Marcia Hobbs, Jonathan Kipnis, and Igor Smirnov for reagents and strains. We thank Sanjay Ram for strains and thoughtful discussions of the work. All experiments with human neutrophils were conducted in accordance with a protocol approved by the UVA Institutional Review Board for Health Sciences Research (IRB HSR #13909).

